# Including measures of high gamma power can improve the decoding of natural speech from EEG

**DOI:** 10.1101/785881

**Authors:** Shyanthony R. Synigal, Emily S. Teoh, Edmund C. Lalor

## Abstract

The human auditory system is adept at extracting information from speech in both single-speaker and multi-speaker situations. This involves neural processing at the rapid temporal scales seen in natural speech. Non-invasive brain imaging (electro-/magnetoencephalography [EEG/MEG]) signatures of such processing have shown that the phase of neural activity below 16 Hz tracks the dynamics of speech, whereas invasive brain imaging (electrocorticography [ECoG]) has shown that such rapid processing is even more strongly reflected in the power of neural activity at high frequencies (around 70-150 Hz; known as high gamma). The aim of this study was to determine if high gamma power in scalp recorded EEG carries useful stimulus-related information, despite its reputation for having a poor signal to noise ratio. Furthermore, we aimed to assess whether any such information might be complementary to that reflected in well-established low frequency EEG indices of speech processing. We used linear regression to investigate speech envelope and attention decoding in EEG at low frequencies, in high gamma power, and in both signals combined. While low frequency speech tracking was evident for almost all subjects as expected, high gamma power also showed robust speech tracking in a minority of subjects. This same pattern was true for attention decoding using a separate group of subjects who undertook a cocktail party attention experiment. For the subjects who showed speech tracking in high gamma power, the spatiotemporal characteristics of that high gamma tracking differed from that of low-frequency EEG. Furthermore, combining the two neural measures led to improved measures of speech tracking for several subjects. Overall, this indicates that high gamma power EEG can carry useful information regarding speech processing and attentional selection in some subjects and combining it with low frequency EEG can improve the mapping between natural speech and the resulting neural responses.

## INTRODUCTION

Scalp-recorded electroencephalography (EEG) provides a non-invasive means of investigating cortical activity with high temporal resolution. This makes it particularly suited for studying neural processes such as speech perception – where humans rapidly convert mechanical vibrations of the air into meaning. In terms of speech, the slow varying acoustic envelope of continuous natural speech was found to be reflected in EEG (Luo and Poeppel 2007; Lalor and Foxe 2010) which is valuable because speech modulations in the 4-16 Hz range have been shown to contain the most important information regarding intelligibility (Drullman, Festen, and Plomp 1994). As a result, many studies tend to focus their analysis around this frequency range when using EEG or magnetoencephalography (MEG) to investigate cortical tracking of the speech envelope (Ahissar et al. 2001; Aiken and Picton 2008; Peelle and Davis 2012; Di Liberto, O’Sullivan, and Lalor 2015).

In contrast to the emphasis on lower-frequency bands in EEG speech research, studies that employ electrocorticography (ECoG) often look at signals in the high gamma range (∼70-150 Hz). High gamma ECoG has also been shown to track the speech envelope (Pasley et al. 2012; Kubanek et al. 2013), even though high gamma and low frequency (LF) activity are thought to result from distinct physiological mechanisms (Edwards et al. 2009). The fidelity of speech tracking in high gamma power (HGP) ECoG data is so high, that most ECoG studies focus exclusively on that frequency range and ignore the data at lower frequencies. Meanwhile, the high frequency content of scalp-recorded EEG is typically disregarded because it is low pass filtered by the skull (Pfurtscheller and Cooper 1975) and smeared by the dura and cerebrospinal fluid (Light et al. 2010), thus resulting in a low signal-to-noise ratio. Nevertheless, we questioned whether there is still useful stimulus-related information dissociable from low frequency data that could be retrieved from high gamma EEG. If so, high gamma EEG could serve as a useful measure for studying speech and language processing in various populations.

In this study, we first investigated the encoding of the temporal speech envelope in HGP EEG. EEG data were recorded as subjects listened to continuous natural speech, and we mapped the EEG (filtered into LF, HGP, and both signals combined) to the temporal speech envelope using linear regression. In a second data set, we investigated whether the inclusion of HGP can improve auditory attention decoding. It is established that envelope tracking in LF EEG is modulated by attention (Kerlin, Shahin, and Miller 2010; Power et al. 2012), and cortical HGP has also been shown to be strongly modulated by attention (Mesgarani and Chang 2012; Zion Golumbic et al. 2013; Dijkstra et al. 2015). Here, we employed a framework that has been successful in ascertaining attentional selection within the context of a task in which subjects attend to one of two concurrent talkers (O’Sullivan et al. 2015). We compared how well attentional selection can be decoded from EEG when using LF, HGP, and a combination of the two. In doing so, we found that for a minority of subjects, the speech envelope and attention are reflected in HGP EEG in a way that is complementary to the information available in LF EEG.

## MATERIALS AND METHODS

### Subjects

Two experimental paradigms were explored in the present study using data from three previously published studies. The first paradigm involved subjects listening to a single speaker and the second involved subjects attending to one of two concurrently presented speakers. Data used in the single speaker paradigm originated from two previous studies examining phoneme level processing and semantic dissimilarity (Di Liberto, Crosse, and Lalor 2018; Broderick et al. 2018). 17 subjects (min = 19 years, max = 31 years, 12 male) were used in total. These studies were approved by the Ethics Committees of the School of Psychology at Trinity College Dublin and the Health Sciences Faculty at Trinity College Dublin. Data used in the attentional selection or cocktail party paradigm was from the control condition of a study which investigated the decoding of auditory attention (Teoh and Lalor 2019). 14 subjects (min = 19 years, max = 30 years, 5 male) took part in the experiment. This study was approved by the Research Subjects Review Board at the University of Rochester. All subjects provided written informed consent and reported no history of hearing impairment or neurological disorders.

### Stimuli and procedure

The single speaker experiment consisted of subjects listening to 20 – 29 trials (approximately 180 s in length) of a mid-20^th^ century audiobook read by one American male speaker (Hemingway 1952). The subjects were tested in a dark, sound-attenuated room and were instructed to attend to a fixation cross in the center of a screen. The storyline was preserved in the trials, with no repetitions or discontinuities present. In the multi-speaker experiment, subjects undertook 20 trials (approximately 60 s in length). They were presented with two stories (Doyle 1902; 1892) simultaneously, narrated by a male and a female speaker. The two audio streams were filtered using head-related transfer functions to simulate spatial separation of the speakers (one speaker at 90 degrees to the left and the other at 90 degrees to the right). The subjects were instructed to attend to the male speaker in all trials, and to minimize motor movements by fixating to a cross at the center of a screen. Subjects then answered four multiple-choice questions on the attended and unattended stories after each trial (which were not analyzed in this work). In all experiments, the stimuli were presented through Sennheiser HD650 headphones at a 44.1 kHz sampling rate using Presentation software from Neurobehavioral Systems (http://www.neurobs.com).

### Data acquisition and Preprocessing

For both experiments, EEG data were acquired at a 512 Hz sampling rate with the BioSemi Active Two system using 128 scalp electrodes (plus 2 mastoid channels that were not analyzed in this work). Each subject’s scalp data were re-referenced to the common average. Noisy channels were determined based on three of EEGLAB’s artifact rejection methods (kurtosis, spectral estimates, and probability distribution; Delorme and Makeig 2004), and spline interpolation was used to reject and recalculate the data on those channels. The EEG was then filtered into two separate bands. The first band (low frequency band) was high pass filtered at 1 Hz and then low pass filtered at 15 Hz using a zero-phase type 2 Chebyshev filter. For the second band, the raw data were bandpass filtered in the high gamma range from 70 – 150 Hz using a 200^th^ order zero-phase FIR filter with a hamming window. The absolute value of the Hilbert transform was taken from the high gamma EEG, and the resulting data was low-pass filtered at 30 Hz to smoothen out any sharp edges that remained in the envelope. Finally, all data were down-sampled to 128 Hz.

### Data analysis

Our analyses for both experiments were based on assessing how strongly the speech signal was represented in our different EEG bands by reconstructing an estimate of the speech envelope from the neural data (Crosse et al. 2016). Speech envelopes were extracted from the stimuli using a gammachirp auditory filterbank which mimics the filtering properties of the human cochlea (Irino and Patterson 1997). Afterwards, the envelopes were normalized between 0 and 1, and the EEG data was z-scored. A backward model (decoder) was employed to reconstruct the speech envelope, *s(t)*, from the neural response, *r(t,n)*, while the decoder, *g(τ,n)*, acted as a linear map between the two. The transformation can be expressed as:

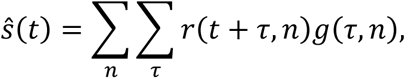

where *ŝ(t)* is the reconstructed speech envelope. The decoder integrates the EEG (with *n* electrodes) over a range of time lags, *τ*, from 0 – 250 ms, the range where low level speech features cause notable EEG responses to occur (Di Liberto, O’Sullivan, and Lalor 2015). The decoder (Figure 1A) was computed by the following operation:

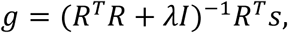

where *R* is the lagged time series, *T*, of the EEG data, λ is the regularization parameter, *I* is the identity matrix, and *s* is the speech envelope.

**Figure 1.**
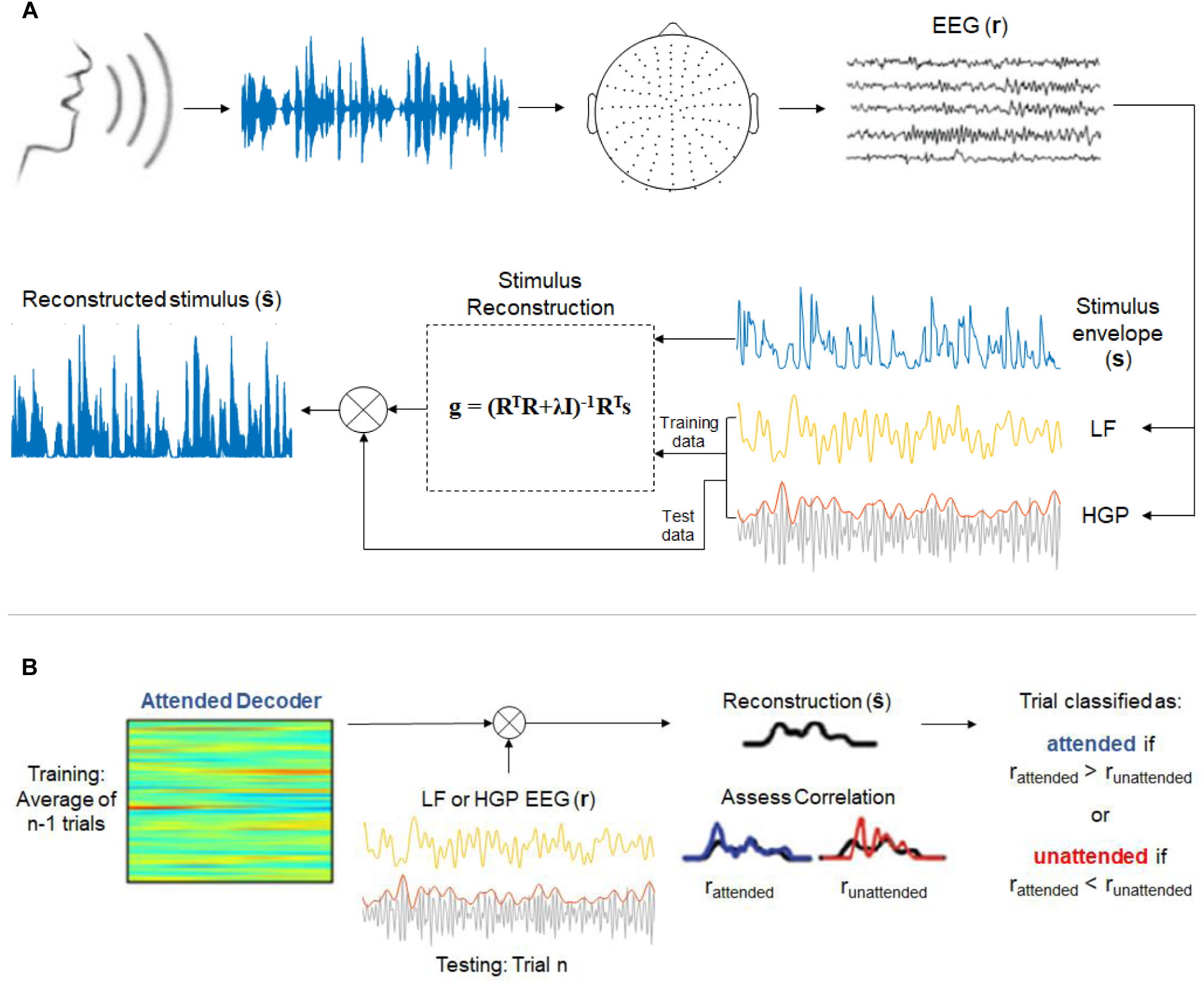
(A) Envelope reconstruction methods (adapted from Di Liberto, O’Sullivan, and Lalor 2015 and Crosse et al. 2016). 128-channel EEG data was collected while subjects listened to continuous, natural speech from a male speaker. Stimulus reconstruction (backward modeling) was used to decode the speech envelope from low frequency (LF, 1-15 Hz) and high gamma power (HGP, 70-150 Hz) EEG recordings. (B) Decoding attention methods (adapted from Teoh and Lalor 2019 and O’Sullivan et al. 2015). Attended decoders (for LF and HGP) were made to reconstruct the attended stimuli. The correlation between the reconstructed stimulus and the attended and unattended speech envelopes were assessed.

Model performance was assessed according to the accuracy in which the speech envelope could be reconstructed using leave-one-out cross-validation. This regression allowed for an optimal regularization parameter to be chosen without overfitting to the training data. The regularization parameter that produced the highest Person’s correlation coefficient between the reconstructed envelope and the actual speech envelope across trials was chosen as the optimal value. Separate decoders were created for the LF and HGP groups. A model was also calculated for the combination of low frequency and high gamma power (LF+HGP) signals, by concatenating the two signals (each 128-channels by delays) to form one matrix of 256-channels by delays.

To decode attention, we employed a framework introduced by O’Sullivan et al. (2015) (Figure 1B). Decoder models that mapped from the EEG data to the speech envelope of the attended speaker were computed for each subject and each trial. The regularization parameter was once again determined based on leave-one-out cross validation. We could then reconstruct the stimulus envelope of a particular trial, *n*, using the average attended decoder of *n-1* trials. The Pearson’s correlation coefficient, r, was computed between the reconstructed envelope and both the actual attended and unattended stimulus envelopes. A trial was deemed correctly classified if the reconstructed envelope was more correlated with the attended envelope rather than the unattended envelope (r_attended_> r_unattended_).

Kruskal-Wallis tests were used to compare stimulus reconstructions based on the different frequency bands (LF, HGP, and LF+HGP). The resulting statistics from the Kruskal-Wallis tests were used to conduct Fisher’s Least Significant Difference (LSD) multiple comparisons tests to determine significance within subjects and between groups. A Wilcoxon Signed Rank test was used when comparing reconstructions between LF and HGP bands only. Testing against chance was completed using permutation tests. In the envelope reconstruction analysis, a null distribution of 1,000 Pearson’s r values was created by finding the correlation between randomly permuted trials of predicted audio envelopes and actual audio envelopes. The true mean correlation coefficient served as the observed value of the test statistic. In the decoding attention analysis, we performed 10,000 permutations to create the null distribution, where for each trial of each permutation we randomly selected a correlation value from either r_attended_ or r_unattended_ to be assigned to bin A, and the other to bin B. The observed value of the test statistic was the percentage of trials where r_attended_ > r_unattended_. The threshold for significance and for above chance performance was p = 0.05 for each test.

## RESULTS

### Though generally weaker than LF EEG, HGP EEG consistently tracks the speech envelope

We first tested how well the speech envelope is reflected in LF (1-15 Hz), HGP (power in the 70-150 Hz range), and LF+HGP (combination of the two) neural signals. To do so, a decoder model was first calculated for each condition. Pearson’s r was then used to quantify the relationship between the actual speech envelope and the reconstructed speech envelope. The grand average reconstruction accuracy (Pearson’s r) for each condition was significantly larger than chance (p = 0.001, permutation test). Next, Kruskal-Wallis tests were used to assess the differences between each decoder. Both the LF and LF+HGP decoders performed significantly better than the HGP decoder (p = 5.8646e-04, p = 0.0047). The LF and LF+HGP decoders, however, displayed no difference in performance (p = 0.5409; Figure 2A). Thus, on a group level, HGP EEG seemed to provide very little additional information regarding the speech envelope.

**Figure 2.**
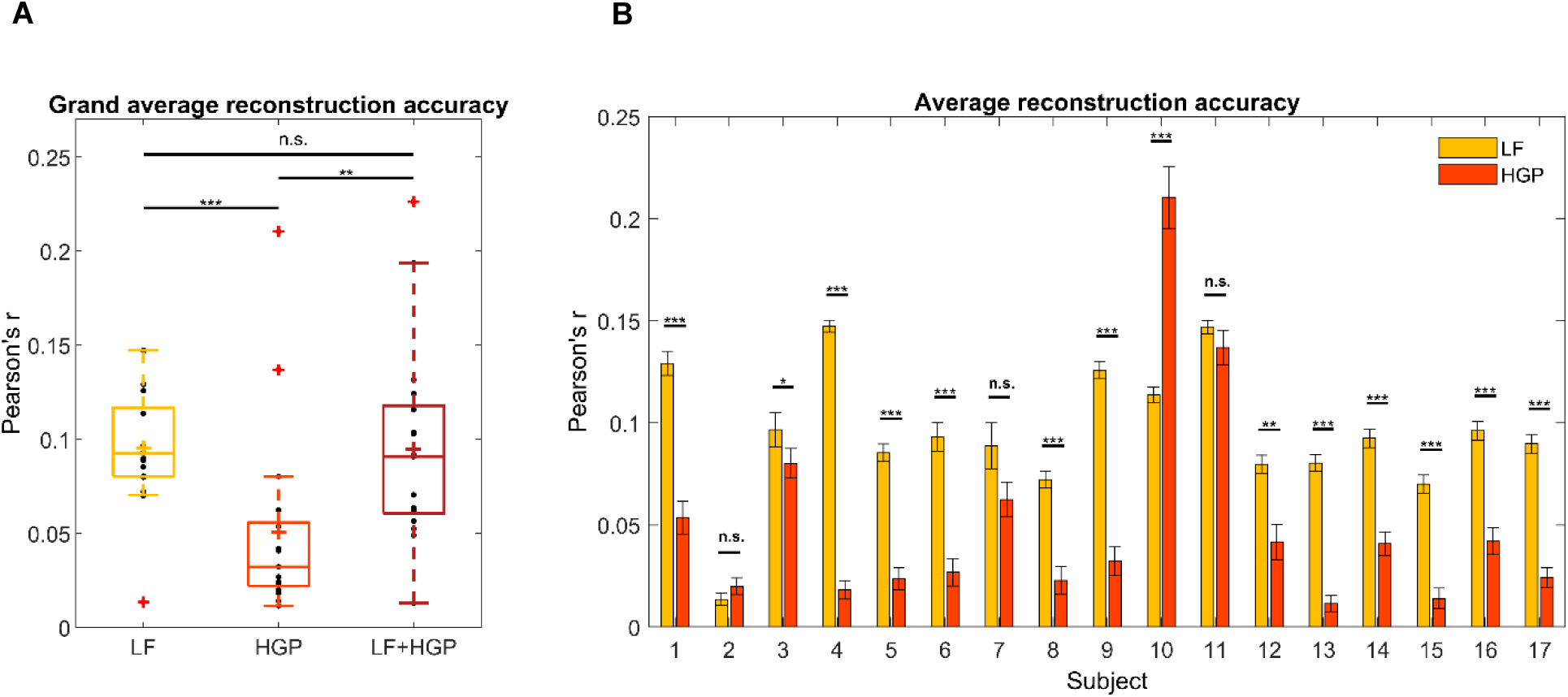
(A) Grand average reconstruction accuracies across trials and subjects for the LF, HGP, and LF+HGP decoders. Each point represents a subject and the colored crosshairs are the mean reconstruction accuracies. Significance is indicated as *** if p ≤ 0.001 using a Kruskal-Wallis test and Fisher’s LSD multiple comparisons test. (B) Mean (± SEM) reconstruction accuracy for each subject across trials for the LF and HGP conditions. Significance is indicated by * if p ≤ 0.05, ** if p ≤ 0.01, and *** if p ≤ 0.001 using a Wilcoxon Signed Rank test.

Since recorded brain activity may vary across individuals due to anatomical differences, we wanted to examine how the LF and HGP decoders performed on a single-subject level. When tested against chance, LF and HGP were significant for all subjects (p ≤ 0.001, permutation test). As expected, the LF decoder worked best for most participants (N = 11, p ≤ 0.001; subject 3, p = 0.0479; subject 12, p = 0.0036; Figure 2B) as shown by the Wilcoxon Signed Rank tests. Surprisingly, there was no difference in reconstruction accuracy for subjects 2, 7, and 11 (p = 0.1943, 0.1072, 0.4115), and the HGP decoder worked better for subject 10 (p = 1.6286e-04). Though uncommon in EEG studies, HGP was able to track the speech envelope comparably or better than LF in some of our subjects.

### Stimulus reconstruction suggests that LF and HGP EEG carry complementary information

Next, we tested if LF and HGP EEG carry complementary information regarding the speech envelope in subjects with comparable HGP measures. To do so, we once again created a combination model (LF+HGP) using 256 channels in total (128 LF EEG channels + 128 HGP EEG channels). The LF+HGP decoder had a higher mean reconstruction accuracy than the LF and HGP decoders alone for three out of seventeen subjects (Figure 3). However, the increase in LF+HGP reconstruction accuracy was only significant in comparison to subject 3’s HGP decoder (p = 0.0025, Kruskal-Wallis test), subject 10’s LF decoder (p = 1.6733e-06), and subject 11’s LF and HGP decoders (p = 2.0059e-05, p =2.0029e-06). This suggests that for these three subjects, HGP and LF EEG carry complementary information.

**Figure 3.**
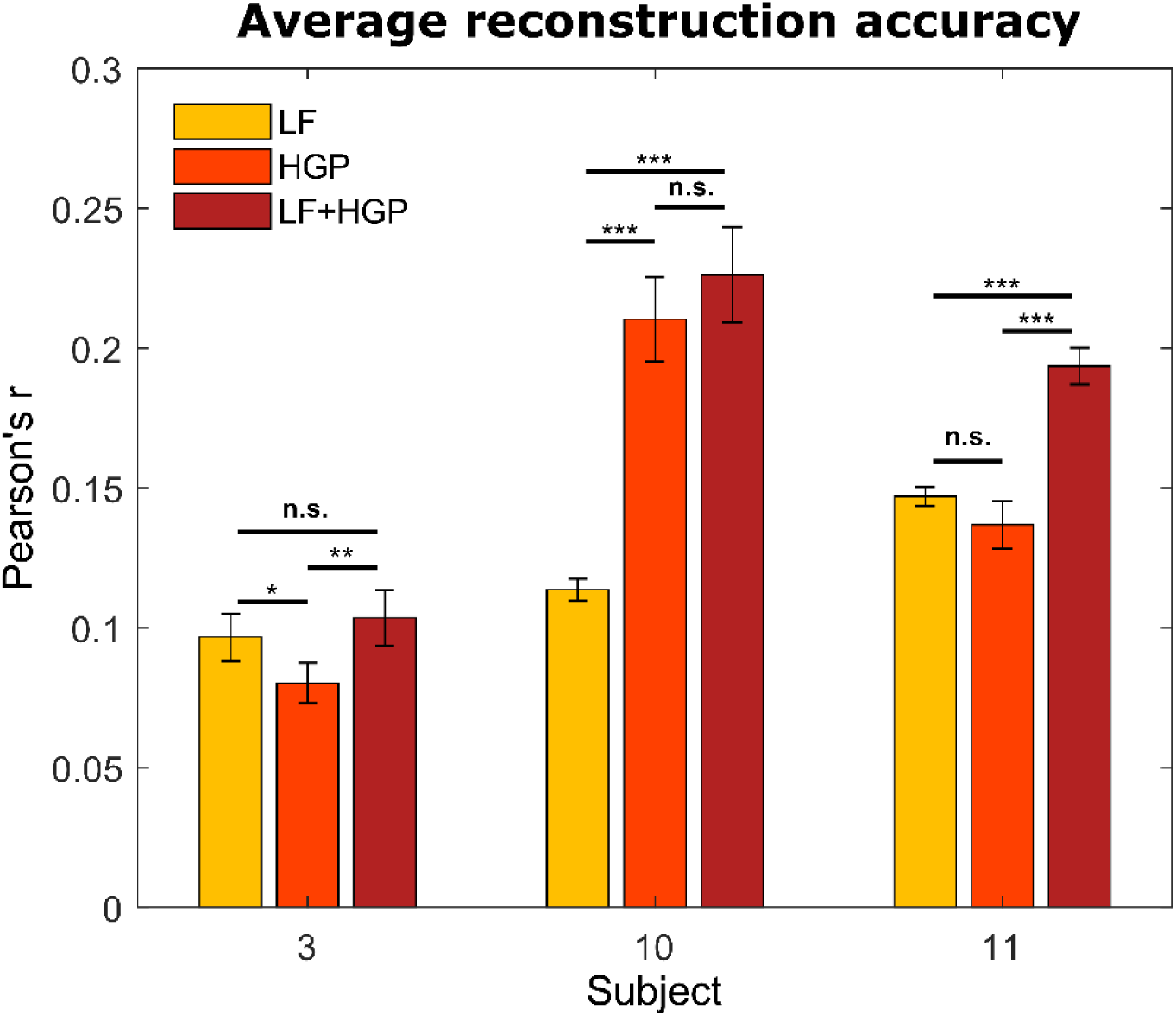
The mean (± SEM) reconstruction accuracies for subjects 3, 10, and 11 using the LF, HGP, and LF+HGP models. Significance is indicated by * if p ≤ 0.05, ** if p ≤ 0.01, and *** if p ≤ 0.001 using a Kruskal-Wallis test and Fisher’s LSD multiple comparisons test.

### LF and HGP responses exhibit different spatiotemporal characteristics to speech

Our stimulus reconstruction analysis suggests that HGP and LF EEG may carry complementary information. To investigate this further, we wanted to examine if there was any evidence that our HGP and LF responses may be derived from different neural generators. To do this we focused on the three subjects who showed robust HGP responses and the distribution of decoder weights across the scalp for their LF and HGP decoders. However, as decoder channel weights cannot be interpreted neurophysiologically (Haufe et al. 2014), we transformed the weights into the forward modeling space using Haufe et al.’s inversion procedure.

With this information, we compared the spatiotemporal profile of the LF and HGP EEG activity for subjects 3, 10, and 11. The left panel of Figure 4A-C depicts spatial dynamics of the forward transformed models (temporal response function or TRF) at various time lags. The LF TRF topographies for the three subjects appeared fairly typical (Crosse et al. 2016) as they alternated in positivity and negativity across time on the frontocentral area of the scalp. On the other hand, their HGP activity displayed a strikingly different distribution with no prominent focus over frontocentral scalp. These results suggest the possibility that the LF and HGP signals we see may have non-identical neural generators and further supports the idea that they may carry complementary information.

**Figure 4.**
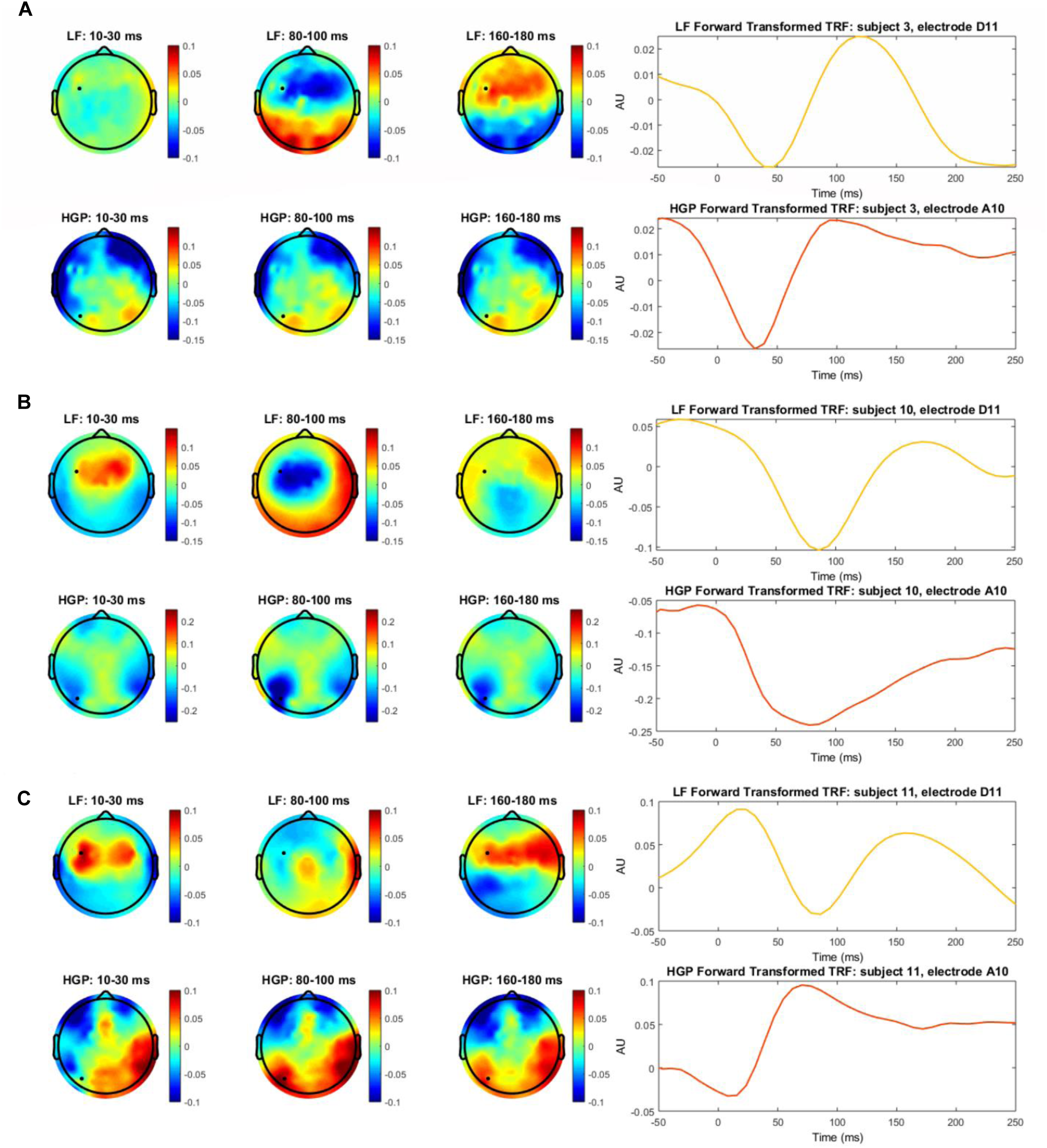
(A) The left panel depicts the transformed decoder weights at each electrode for subject 3. The top three topographies were calculated using LF EEG and the bottom three using HGP EEG. The black marker indicates the location for electrodes D11 and A10. The right panel shows subject 3’s temporal response function (TRF) calculated by transforming the decoder weights into the forward model domain. The LF TRF (yellow) is plotted for electrode D11 and the HGP TRF (red-orange) is plotted for electrode D31. (B, C) Same information as (A), but shown for subjects 10 and 11 respectively.

TRFs provide an example of how the speech envelope is transformed into neural responses over time at specific electrodes. The right panel of Figure 4 shows the forward transformed TRFs for subject 3, 10, and 11 at the electrode indicated in their topographies. The TRFs at the given electrodes are highly correlated for subjects 3 and 10 (r = 0.7985, r = 0.8454, Spearman’s correlation) and moderately correlated for subject 11 (r = −0.6762). While there are differences in the time course between LF and HGP TRFs – again supporting the notion that the neural generators might differ – the general timing of the two TRFs is similar for these three subjects supporting the notion that the HGP TRFs we see in these subjects is capturing real responses to our speech stimuli.

### HGP also improves the decoding of auditory attention in some subjects

It has previously been shown that the LF envelope tracking response is modulated by attention (Ding and Simon 2012), and that single trial data from a task in which subjects attend to one of two concurrent talkers can be decoded to ascertain attentional selection (O’Sullivan et al. 2015). Given our finding that HGP contains informative temporal envelope information for a subset of subjects, we tested whether this signal is similarly modulated by attention and if it could be exploited to improve our ability to decode attention in multi-speaker situations. This was tested on a separate group of subjects (N = 14) from those used in the single speaker paradigm.

The decoding accuracy here represents the percentage of trials in which the reconstructed stimuli were more correlated with the attended stream rather than the unattended stream (r_attended_ > r_unattended_) for LF, HGP, and LF+HGP EEG signals. Exploring this on a group-level showed a similar trend as our initial envelope tracking results from Figure 2A. The decoding accuracy was significantly larger than chance for each model (p < 0.001, permutation test; Figure 5A). The HGP decoder was able to decode auditory attention, but it did not perform as well as the LF and LF+HGP decoders (p = 6.9846e-04, p = 9.2431e-04) once again. LF alone decoded attention better than LF+HGP on average (84.64% vs 83.57%), but this difference was not significant (p = 0.9382). We also examined how well we were able to decode auditory attention for individual subjects and found four whose decoding improved with HGP or LF+HGP EEG signals using permutation tests. Figure 5B shows the decoders that were able to track the temporal dynamics of the attended speaker significantly better than chance (p ≤ 0.05). Subjects 2, 3 and 10 reliably tracked the attended speaker’s speech envelope best using LF+HGP EEG signals (95%, 100%, and 70% respectively), whereas subject 11 reliably tracked the attended speaker best using HGP EEG (70%). Interestingly, subject 10’s LF and HGP signals alone did not decode attention above chance but combining the two significantly improved decoding performance (p = 0.0182).

**Figure 5.**
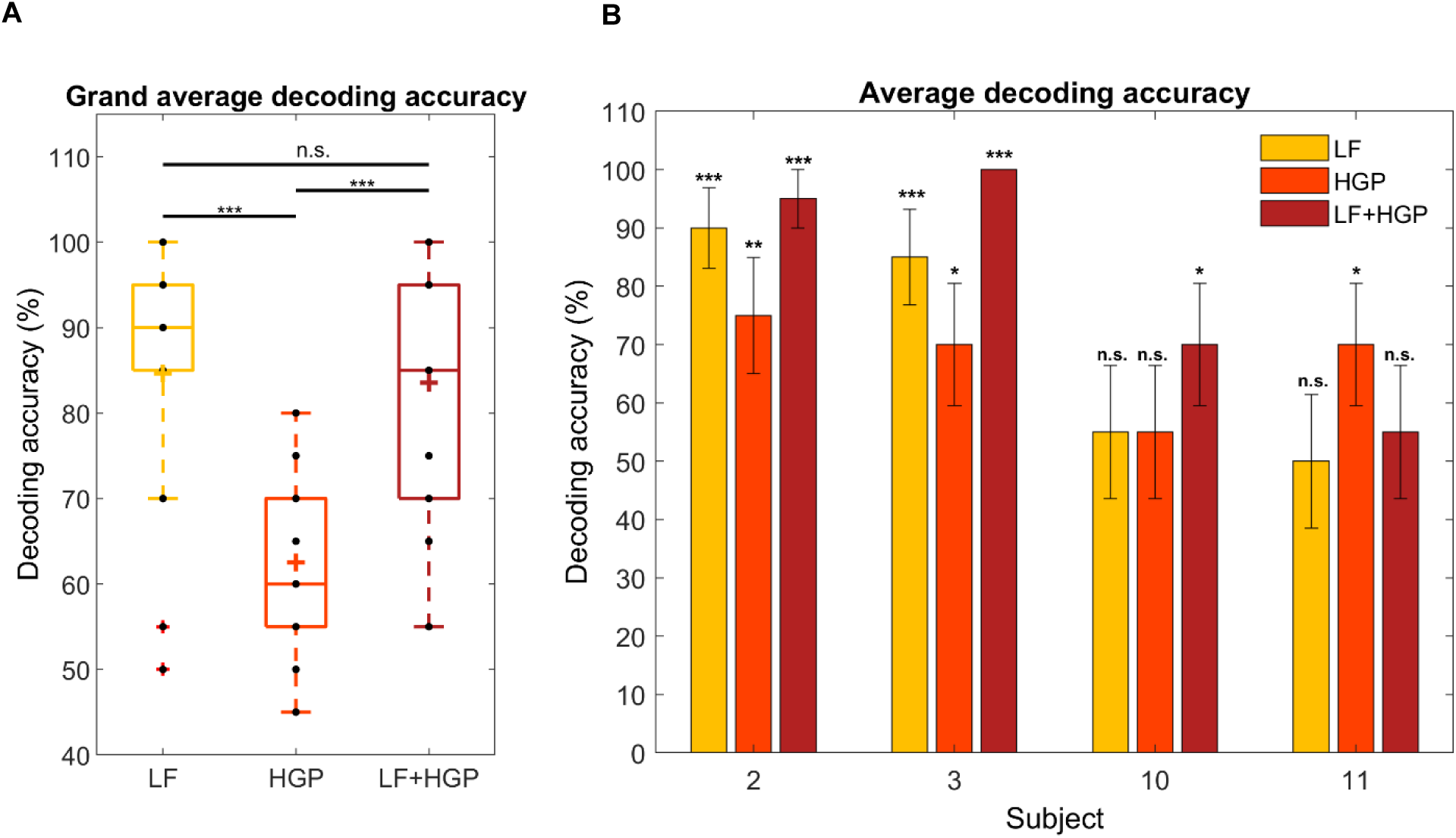
(A) Grand average decoding accuracies across trials and subjects for the LF, HGP, and LF+HGP decoders. Each point represents a subject and the colored crosshairs are the mean accuracies. Significance is indicated as *** if p ≤ 0.001 using a Kruskal-Wallis test and Fisher’s LSD multiple comparisons test. (B) The mean (± SEM) decoding accuracies for subjects 2, 3, 10, and 11 using the LF, HGP, and LF+HGP models. Significance is indicated by * if p ≤ 0.05, ** if p ≤ 0.01, and *** if p ≤ 0.001 using permutation tests.

## DISCUSSION

Our present research investigated the extent to which the speech envelope and attentional selection are reflected in low frequency phase and high gamma power EEG signals. Given the success of using HGP to track the dynamics of speech in ECoG studies, we wanted to examine if any useful high frequency EEG activity remained even after being smeared by brain tissue and filtered by the skull. In this study, linear regression techniques were used to map between neural responses and the acoustic envelope of speech. Our results demonstrate that HGP EEG activity is capable of carrying information regarding the speech envelope and attention complementary to that of LF EEG, as shown by a minority of our subjects.

Previous studies using EEG techniques to study gamma band activity in the human auditory system typically examined activity in the 40 Hz range (Jokeit and Makeig 1994; Krause et al. 1998; Gurtubay et al. 2001; Hald, Bastiaansen, and Hagoort 2006). This could be due to EEG’s poor signal-to-noise ratio (Crone et al. 2001), poor spatial resolution, or sensitivity to muscle artifacts (Llorens et al. 2011). Another possible reason is that HGP is generated by highly focal sources (Jerbi et al. 2009) which in turn produce lower amplitudes on the scalp (Nunez and Srinivasan 2006; Jerbi et al. 2009). Nevertheless, EEG provides valuable temporal information regarding electrical activity that relates to various neural events.

We have shown that HGP, though typically weaker than LF EEG, can consistently track the temporal dynamics of speech. In some subjects, HGP EEG alone could decode the speech envelope and auditory attention best, and in others, combining LF and HGP neural signals further improved decoding. These results suggest that LF and HGP EEG carry complementary information, in support of Belitski et al. who used measures of synergy to show that there is very little redundant information carried between low frequency and high frequency neural signals (Belitski et al. 2010).

Our results are also in line with Akbari et al. who found that combining low frequency and high gamma ECoG signals better reconstructed the speech envelope for their BCI system (Akbari et al. 2019). Similarly, a cocktail party attention study showed that both LF phase and HGP ECoG signals can track the envelope of the attended speaker, and combining the two may optimize attention encoding (Zion Golumbic et al. 2013). Each of these studies provide further support for interactions between HGP and LF phase neurophysiological signals during sensory processing (Bruns and Eckhorn 2004; R. T. Canolty et al. 2006; Osipova, Hermes, and Jensen 2008; Voytek et al. 2010) and for the notion that speech encoding may involve the combination of the two (Nourski et al. 2009; Zion Golumbic et al. 2013).

The TRF weightings of subjects in whom we were able to detect high gamma electrophysiology displayed different spatial patterns for LF and HGP responses. HGP and LF signals are said to originate from different locations in the brain (Crone et al. 2001; Edwards et al. 2009), supporting our findings in the left panel of Figure 4. Studies also suggest that HGP is mainly localized to the superior temporal gyrus (Crone et al. 2001; Towle et al. 2008; Sinai et al. 2009) in contrast to LF activity which is more spatially distributed across temporal and some frontal and parietal regions of the brain (Crone, Sinai, and Korzeniewska 2006; Ryan T. Canolty et al. 2007; Zion Golumbic et al. 2013). Although neural signals are generally spatially smeared in EEG measures, we still saw differences in the scalp patterns which may be indicative of different sources of LF and HGP activity. Of course, we need to be somewhat circumspect here, because our contrasting scalp patterns may have only arisen by virtue of differences in the biophysics of how signals at high and low frequencies project to the scalp (Buzsáki, Anastassiou, and Koch 2012). Indeed, the broadly similar timing of our LF and HGP TRFs (Figure 4) might support that notion. The abovementioned ECoG work, however, ultimately supports the idea of different generators.

Gamma activity assessed through non-invasive means has been shown to play a role in a variety of neural processes such as working memory (Tallon-Baudry et al. 1998; Howard et al. 2003; Mainy et al. 2007; Roux and Uhlhaas 2014), motor and sensorimotor function (Medendorp et al. 2007; Ball et al. 2008; Cheyne et al. 2008), and visual processing (Adjamian et al. 2004; Hoogenboom et al. 2006; Fründ et al. 2007). Here we show that HGP also has value when studying speech processing and auditory selective attention – albeit in a minority of subjects. In these subjects, high gamma activity supplemented lower frequencies to increase the sensitivity to speech and attention related processes. Therefore, it is worth investigating HGP in all subjects as this increase in sensitivity could be beneficial, for instance, for the use of future EEG-enabled hearing devices.

## AUTHOR CONTRIBUTIONS

EST collected the cocktail party data and SRS analyzed the data. SRS, EST, and ECL interpreted the data and wrote the article.

